# Activity of epigenetic inhibitors against *Babesia divergens*

**DOI:** 10.1101/2020.12.04.411777

**Authors:** Leen N. Vanheer, Björn F.C. Kafsack

**Affiliations:** Department of Microbiology & Immunology, Weill Cornell Medicine, New York, NY, USA

## Abstract

Babesiosis in a tick-borne parasitic disease of humans and livestock, that has dramatically increased in frequency and geographical range over the past few decades. Infection of cattle often causes large economic losses, and human infection can be fatal in immunocompromised patients. Unlike for malaria, another disease caused by hemoprotozoan parasites, limited treatment options exist for Babesia infections. As epigenetic regulation is a promising target for new anti-parasitic drugs, we screened 324 epigenetic inhibitors against *Babesia divergens* blood stages and identified 75 (23%) and 17 (5%) compounds that displayed ≥90% inhibition at 10 µM and 1 µM, respectively, including over a dozen compounds with activity in the low nanomolar range. We observed differential activity of some inhibitor classes against *Babesia divergens* and *Plasmodium falciparum* parasites and identified pairs of compounds with a high difference in activity, despite a high similarity in chemical structure, highlighting new insights into the development of epigenetic inhibitors as anti-parasitic drugs.

## INTRODUCTION

Babesiosis is an emerging parasitic disease caused by the intra-erythrocytic *Babesia* parasite and has many clinical features similar to malaria infection. Babesia infections in cattle are widespread and lead to economic losses through death, reduction in meat and milk yield, and the cost of control measures. More than 100 *Babesia* species have been identified but only a few infect humans. However, human babesiosis is an increasing concern worldwide, as the number of reported cases have increased over the last decades and the geographical rage of transmission has expanded ^1,2^. In Europe, *Babesia divergens* is responsible for most human babesiosis cases ^3^. In the US, the majority of human *Babesia* infections is caused by *Babesia microti*, although cases of *B. divergens*-like organisms have been reported as well ^4^. Transmission occurs through the bite of an infected tick or occasionally through blood transfusion, which has prompted the screening of the blood supply for Babesia parasites in an increasing number of U.S. states ^5^. Symptomatic human babesiosis is manifested by malaria-like symptoms, such as fever and general malaise. Treatment recommendations for human babesiosis are a combination of atovaquone and azithromycin for mild to moderate babesiosis and clindamycin plus quinine for severe infection ^6^. However, for immunocompromised patients, cases of treatment failure for both regimens have been reported ^7,8^ and babesiosis can lead to organ failure and death. Asplenic patients, in particular, are at high risk for relapsing infections and require long antimicrobial treatment ^9^. Recently, tafenoquine, a newly FDA-approved drug for malaria treatment, showed activity against *Babesia microti* in mice but further studies are needed ^10^. As treatment options for relapsing Babesia infection are limited, research into new drugs for babesiosis is critical.

In eukaryotes, epigenetic regulation of gene expression, mediated by small modifications of nucleosomes and on DNA itself, has been found to be critical for cellular homeostasis and differentiation ^11^. In a recent screen of 324 commercially available epigenetic inhibitors against *Plasmodium falciparum* ^12^, we showed that 54 compounds exhibited ≥50% inhibition at 1 µM *in vitro*, suggesting that the epigenetic machinery could be a promising novel drug target. Since *Plasmodium* and *Babesia* are related parasite species with similarly complex life cycles that share much of the epigenetic regulatory machinery, we decided to determine the activity of these epigenetic inhibitors against *Babesia divergens*, for which an in vitro culture system and drug assays are established.

## METHODS

Commercially available libraries of 324 epigenetic inhibitors from Selleckchem (Houston, TX) and Cayman Chemicals (Ann Arbor, MI) were purchased. Libraries were aliquoted and diluted in DMSO to 2 mM and 0.2 mM in V-bottom 96-well plate and stored at −80°C.

*Babesia divergens* (Bd Rouen 1987 strain ^13^) was grown in vitro in human A+ RBC at low parasitemia. Cultures were flushed with 90% nitrogen, 5% oxygen and 5% carbon dioxide and cultured at 37°C. SYBR Green based growth assays were used to determine in vitro activity of the epigenetic inhibitors against *B. divergens* ^14,15^. Briefly, flat-bottom 96-well plates at a total of 200 μL per well and 0.5% DMSO, at a final 5% hematocrit, 0.5% parasitemia for *B. divergens* were incubated at 37°C for 72 hours and thereafter frozen at −80°C. Upon thawing, 100 μL of SYBR Green (ThermoFisher) diluted in lysis buffer (0.2 μL 10,000X SYBR Green per mL lysis buffer) was added to each well and plates were shaken in the dark at room temperature for 1 hour. Fluorescence was then measured using a Molecular Devices SpectraMax ID5 plate reader. Values were normalized to solvent-treated controls (included in triplicate on each plate). EC50 values were calculated using the nlmLS function of the minpack.lm package (v1.2-1) of the R statistical package (v3.6.0).

Protein sequences of histone modifying enzymes were retrieved from PlasmoDB and PiroplasmaDB ^16,17^. Orthologs were identified using annotated ortholog groups as well as reciprocal BLAST searches. NCBI Conserved Domain Search ^18^ was used to identify conserved protein domains. Protein sequences were aligned using MUSCLE ^19^ and visualized using MView ^20^. In cases where two consecutive genes were identified as orthologs, the geneID for the ortholog containing the catalytic domain was indicated as the ortholog.

Structural feature (SkelSphere) analysis and Activity Cliff Analysis were performed using Osiris DataWarrior v5.2.1, at 80% chemical structure similarity cut-off. Structure-Activity Landscape Index (SALI) values ^21^ were calculated as:

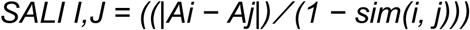

in which *Ai* and *Aj* are the activities of compounds *i* and *j*, and *sim(I,j)* is the similarity coefficient between the two molecules.

## RESULTS AND DISCUSSION

### Evolutionary conservation of epigenetic modifying enzymes in piroplasmid parasites

Among eukaryotes, the most common epigenetic modifications are acetylation of histone lysine residues, methylation of histone lysine and arginine residues, and methylation of deoxycytidine on DNA. We were able to identify orthologs for most of the enzymes classes that place and remove these marks in the genomes of piroplasmid parasites, which includes Babesia and Theileria species (Figure 1, Supplemental Table 1, Supplemental Dataset 1). Orthologs to seven of the eleven <u>S</u>u(var)3-9/<u>E</u>nhancer of Zeste/<u>T</u>rithorax (SET)-domain-containing lysine-specific histone methyltransferases (KMT) present in *P. falciparum* could be identified in at least one piroplasmid genome. Orthologs of PfSET4 and PfSET5 were absent in the entire clade while an ortholog to PfSET6 could only be identified in *B. microti*. Orthologs to PfSET9 could be identified in all piroplasmida genomes but are missing key residues within the catalytic SET domain thus making it unlikely that these have retained methyltransferase activity. Most piroplasmid genomes also retain two additional trithorax-like SET-domain proteins found in other apicomplexan parasites that were lost in malaria parasites. Histone demethylases were notably reduced compared to *P. falciparum*, with *B. divergens* and *B. bovis* only encoding a single member of the Jumonji and LSD demethylase families.

**Figure 1.**
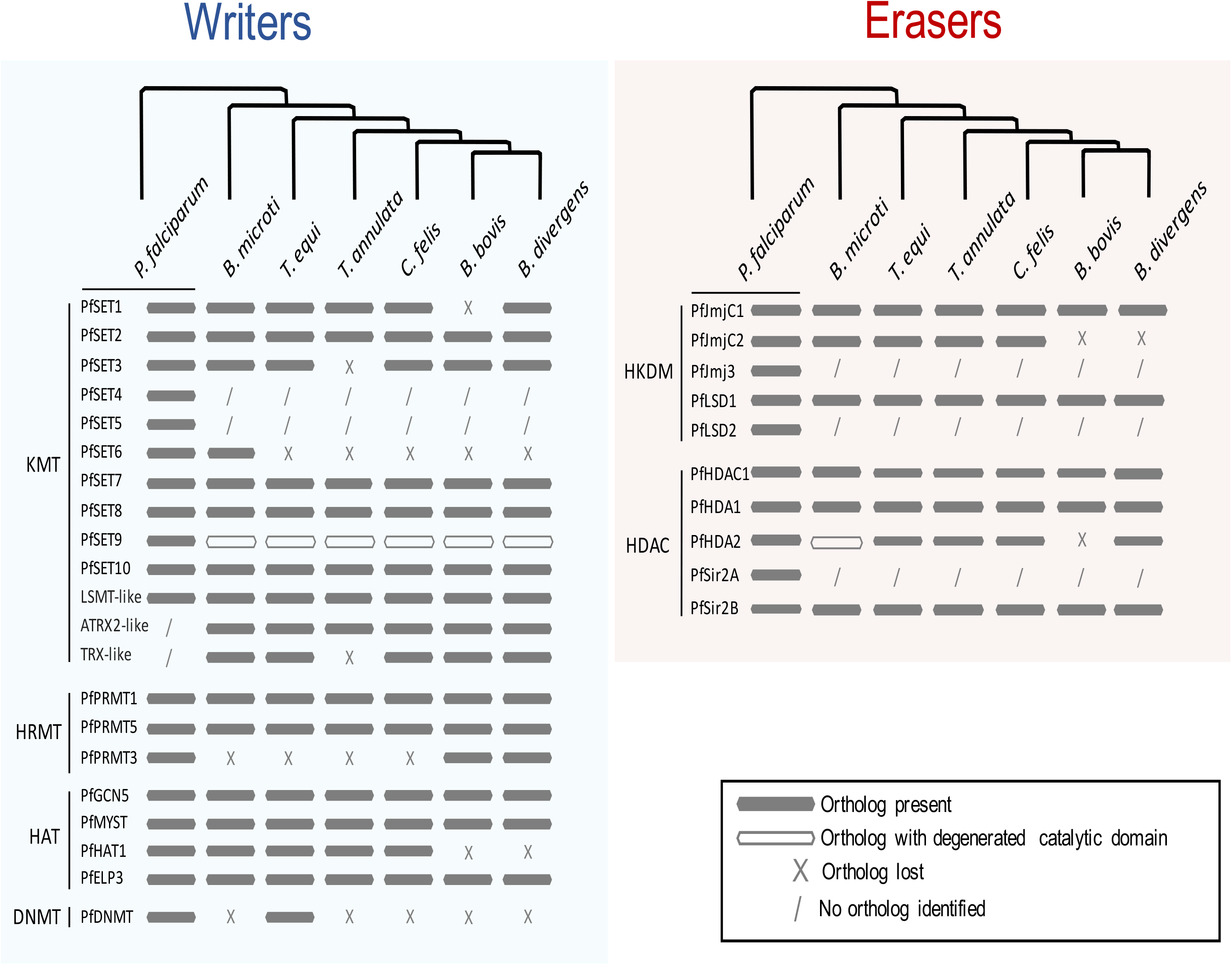
Evolutionary conservation of epigenetic modifying enzymes in piroplasmid parasites. Comparison of epigenetic modifying enzyme orthologs between *P. falciparum, B. microti, T. equi, T. annulata, C. felix, B. bovis* and *B. microti*, with a representation of the evolutionary relationship between these species.

All three the histone arginine methyltransferases (RMTs) in *P. falciparum* had orthologs in *B. divergens*, but the PRTM3 ortholog was lost outside the Babesia *sensu stricto* clade (represented by *B. bovis* and *B. divergens*). Histone acetyltransferases (HATs) and deacetylases (HDACs) were generally conserved between malaria parasites and the piroplasmida. Notable is the absence of Sir2A from all piroplasmid genomes and loss of HAT1 from the Babesia *sensu stricto* clade. An ortholog of a proposed Cytosine-5 DNA methyltransferase in *Plasmodium falciparum* ^22^ is present in *Theileria equi* but could not be identified in the other genomes examined.

### Activity of epigenetic inhibitors against *Babesia divergens*

Given these differences in histone modifying enzymes between *P. falciparum* and the piroplasms, we decided to test the susceptibility of *B. divergens* to a library of epigenetic inhibitors previously screened against P. falciparum ^12^. Of the 324 compounds tested, 125 (39%) showed ≥50% inhibition at 10 µM against *B. divergens* blood stages, of which 46 (14%) retained greater than half-maximal activity at 1 µM (Figure 2A, Supplemental Figure 1 and Table 2, Supplemental Dataset 2). 75 (23%) and 17 (5%) of compounds exhibited greater than 90% inhibition at 10 µM and 1 µM, respectively. Dose-response curves were performed for 17 compounds with sub-micromolar EC90 values (Figure 2B). Of these, the HDAC inhibitors quisinostat and apicidin were the most potent compounds, with EC50 values as low as 5 - 6 nM. These top hits included two FDA-approved drugs mitomycin C and panobinostat with EC50 values of 63 nM and 27 nM, respectively. Peak plasma concentration of panobinostat dosage indicated for treatment of multiple myeloma only correspond to EC75 for Babesia making it an unlikely candidate for treatment ^23,24^. Mitomycin C is a CpG DNA crosslinking agent indicated, indicated for gastric and pancreatic adenocarcinoma treatment and was recently also approved for low-grade upper tract urothelial cancer.^25^. Intravenous administration during chemotherapy leads to plasma concentration of 5 µM ^26^, around 20-fold higher than its EC90 value against *B. divergens*. We previously determined toxicity of selected compounds against human HepG2 cells ^12^. Several compounds displayed only moderate toxicity against HepG2 cells even at 1 µM (Figure 2B). This drug screen identifies promising compounds for additional SAR studies for possible use as a new class of anti-*Babesia* drugs.

**Figure 2.**
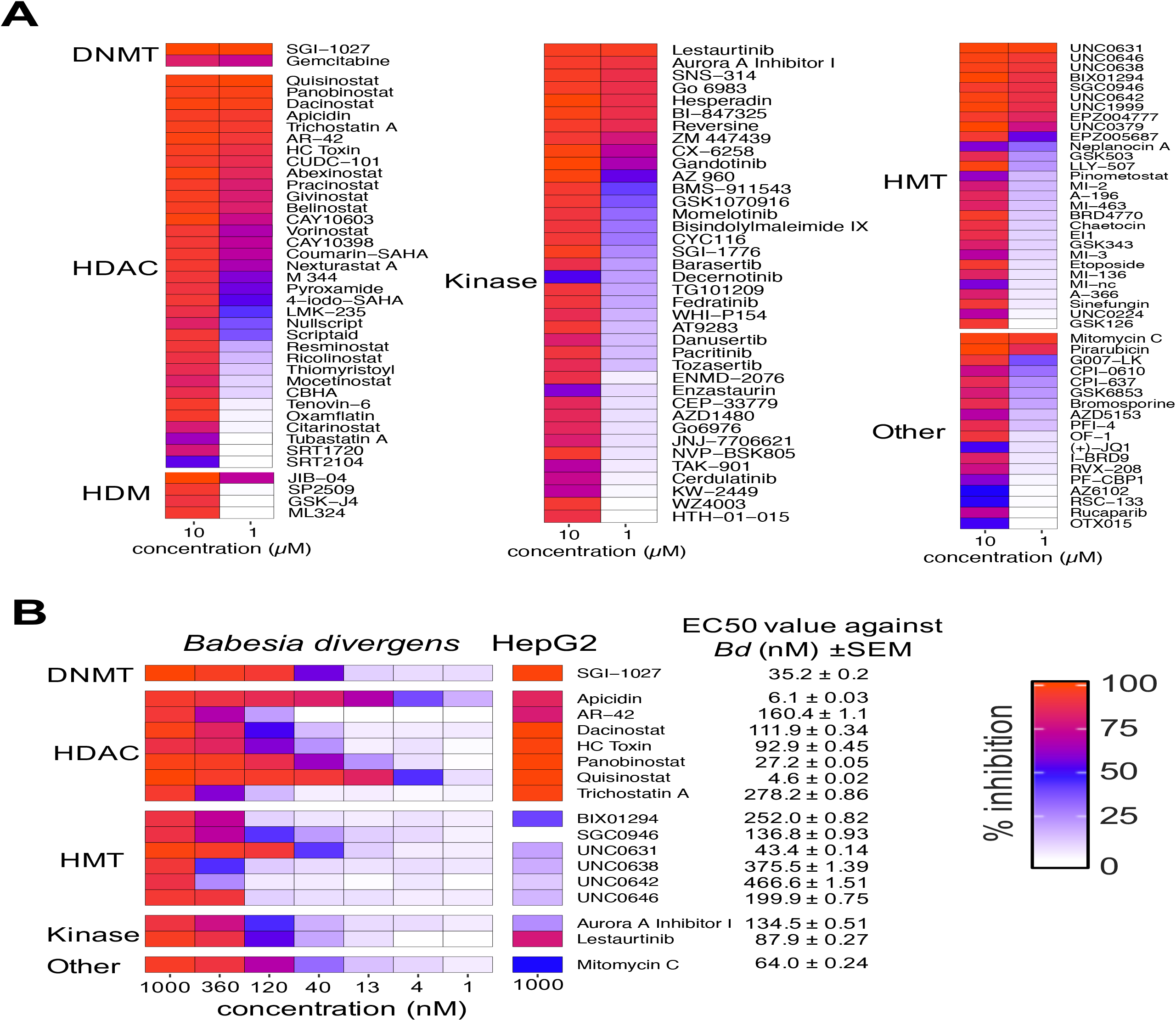
Activity of epigenetic inhibitors against *Babesia divergens*. **(A)** 125 compounds with ≥50% inhibition at 10 µM. Heatmap of mean percent inhibition at 10 and 1 µM compared to solvent-treated controls (n=3). Compounds are grouped based on the reported epigenetic process affected in higher eukaryotes: Histone deacetylation (HDAC), histone acetylation (HAT), histone methylation (HMT), Histone Demethylases (HDM), DNA methylation (DNMT), and “Other”. **(B)** Dose response analysis for 17 compounds with sub-micromolar EC50 values (n=2), with corresponding HepG2 inhibition at 1 µM.

### Differential activity of epigenetic inhibitors against *B. divergens* and *P. falciparum*

Next, we compared these results to our recently completed screen of this library against *P. falciparum* blood stages ^12^. A similar number of compounds had >50% inhibition at 1 µM against both species (46 against *B. divergens* compared to 54 against *P. falciparum*). For both species, compounds targeting histone methylation, deacteylation, demethylation or phosphorylation were the most active, while compounds targeting histone acetylation, PARPylation, histone reader domains, DNA methylation or other pathways had little to no activity (Supplemental Table 1).

Twenty-five compounds with >50% inhibition at 1 µM against one species exhibited greater than two-fold difference in activity in the other (Figure 3). HDAC inhibitors were generally less active against *B. divergens* than against *P. falciparum*, with 18% of the 85 HDAC inhibitors in the library showing ≥90% inhibition at 1 µM against *P. falciparum* versus only 8% for *B. divergens* (Supplemental Table) 2. Interestingly, several HDAC inhibitors with high differential activity target the human HDAC1 (class I) or HDAC6 (class IIb), suggesting that the HDACs may be more divergent from the human enzymes in *B. divergens* than in *P. falciparum*. An additional 25 compounds exhibited greater than 75% inhibition against both *P. falciparum* and *B. divergens*. Thirteen of these were HDAC inhibitors many of which have activity against multiple HDAC classes. Surprisingly, of the 15 HMT inhibitors in Figure 3B, the four compounds with greater activity against *B. divergens* target either ethery the H3K79 HMT DOT1L or the H3K27 HMT EZH1/2 in humans, orthologs to which are absent from both species. As in our previous study, the DNMT inhibitor SGI-1027 was also among the most active compounds against *B. divergens*, despite no identifiable DNMT ortholog in *Babesia* species and PfDNMT being dispensable for asexual growth ^27^, suggesting that SGI-1027 likely has one or more alternative targets.

**Figure 3.**
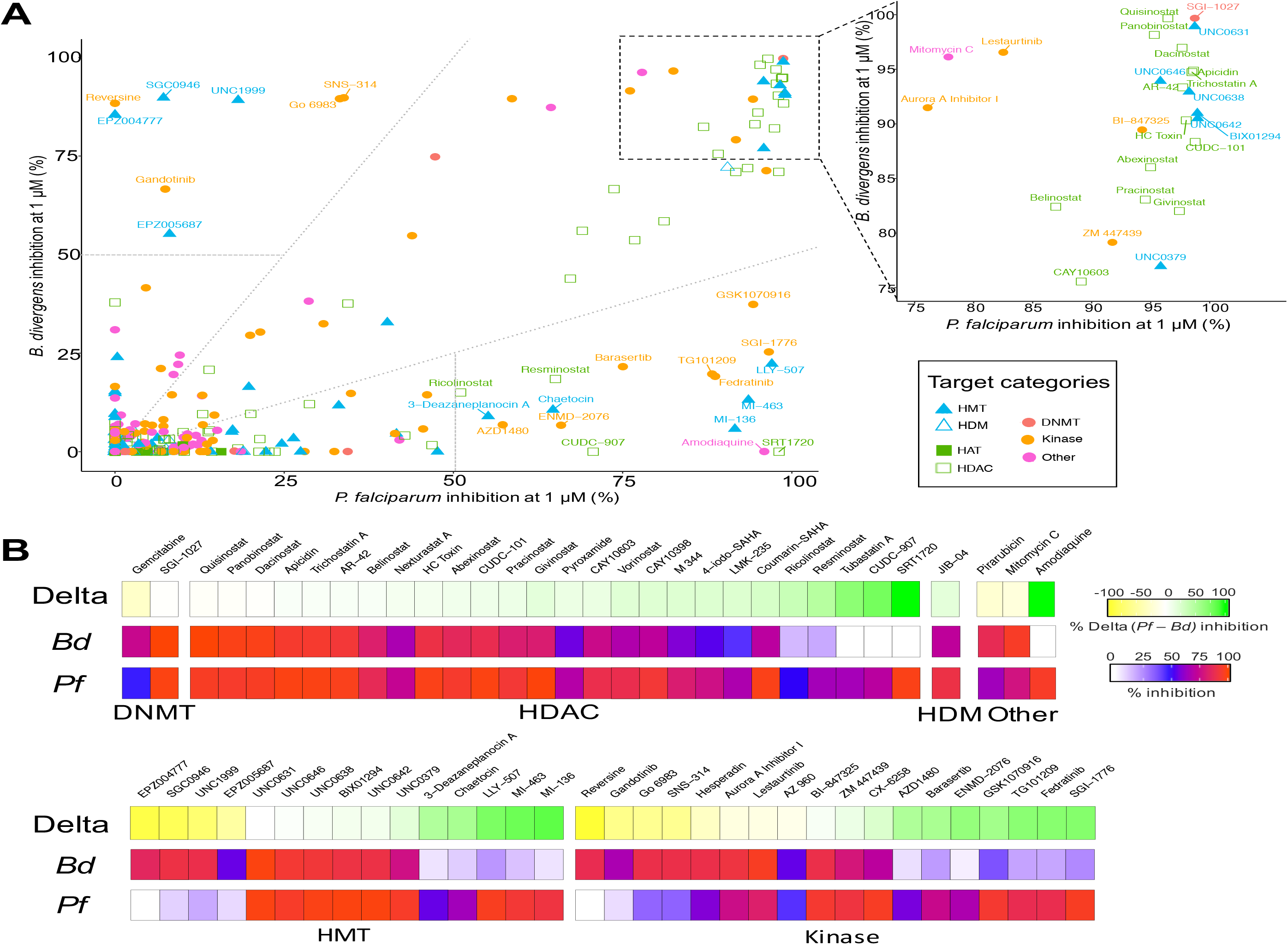
Differential activity of epigenetic inhibitors against *B. divergens* and *P. falciparum*. **(A)** Scatterplot comparing %inhibition at 1 µM against *B. divergens* and *P. falciparum*. Compound names are indicated for compounds with more than 2-fold difference in activity (dotted lines) and more than 50% inhibition at 1 µM against one species (dashed lines). An enlarged scatterplot with labelled compound names is displayed for compounds with ≥75% at 1 µM against both species. **(B)** Heatmap of compounds with at least 50% inhibition at 1 µM against one species, ordered by the delta activity (% *Pf* inhibition - % *Bd* inhibition) and grouped by proposed target category.

### Similarity and activity cliff analysis of activity against *B. divergens* and *P. falciparum*

Structural feature analysis of all 324 unique compounds revealed five clusters of four compounds or more with >80% structural similarity (**Supplemental** Figure 2), including seven HDAC inhibitors with a common hydroxamate-based scaffold, seven HMT inhibitors sharing a common diaminoquinazoline backbone (Figure 5) and three HMT inhibitors an 1H-indazole-4-carboxamide scaffold (Figure 4B). Activity Cliff analysis identifies pairs with high differential activity, despite high structural similarity. Delta activity and SALI values are plotted for all pairs of the library in Supplemental Figure 3. Compound pairs of interest have >50% delta activity and >80% structural similarity (Figure 4A). Twelve pairs were activity cliffs in both *P. falciparum* and *B. divergens*, while three only had more than 50% delta activity in *B. divergens* and 4 only in *P. falciparum*.

**Figure 4.**
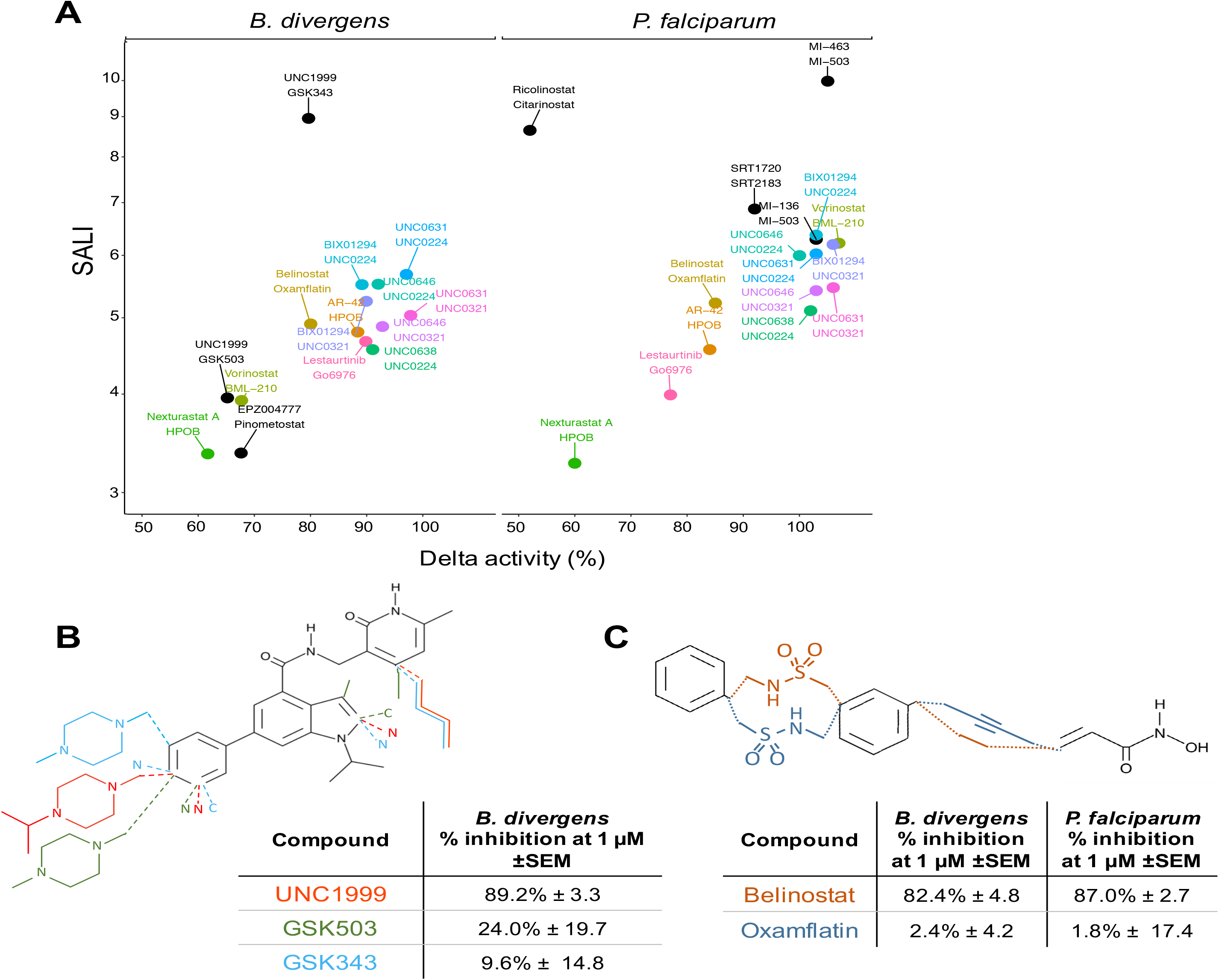
Activity cliff analysis. **(A)** Scatterplot of 19 activity cliff pairs with >50% delta activity and >80% structural similarity, grouped by species. Compound pairs that display an activity cliff in both species are indicated in matching colors. **(B-C)** Examples of activity cliff pairs with respective chemical structures and in vitro activity at 1 µM.

**Figure 5.**
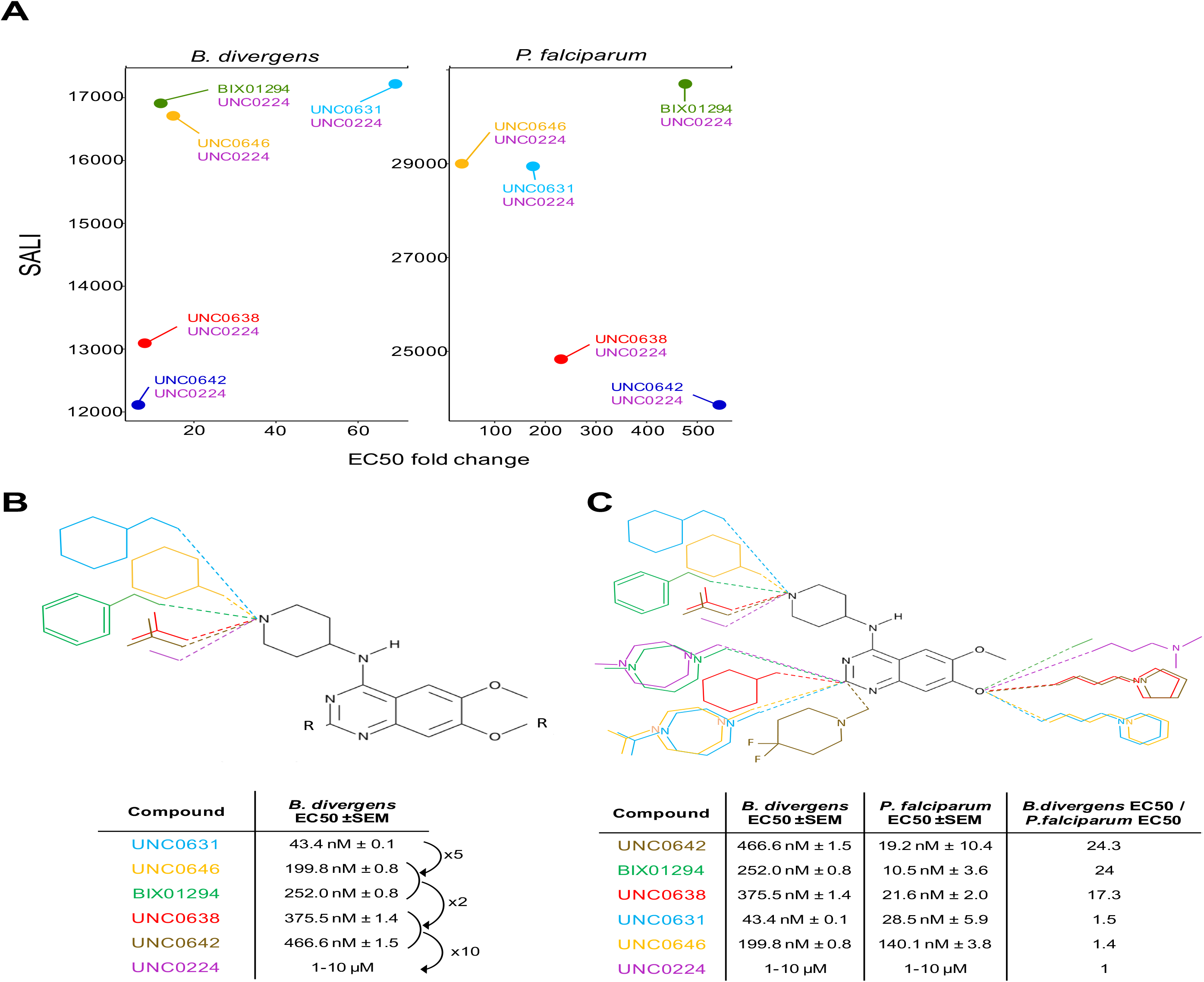
HMT inhibitors with a diaminoquinazoline backbone. **(A)** Pairs of diaminoquinazoline compounds with >60% similarity and >50% delta activity at 1 µM. **(B)** Activity of compounds with a diaminoquinazoline backbone against *Babesia divergens*. **(C)** Changes in chemical structure that confer differential activity against both species.

The HMT inhibitor UNC1999 displays an activity cliff for activity against *B. divergens* when paired with GK343 and GSK503, despite the former having 91% structural similarity (Figure 4B). Interestingly, the IC50 values for mammalian EZH2 enzyme inhibition are similar for all three compounds, while UNC1999 is the only compound with potent EZH1 inhibition as well. Kinase inhibitors belinostat and oxamflatin show activity cliffs for both species (Figure 4C). Additional structural comparisons of the remaining activity cliff pairs can be found in Supplemental Figure 4. These activity cliff pairs provide insight into the structural features that confer activity against *P. falciparum* and *B. divergens*.

### Differential HMT inhibitors with a diaminoquinazoline backbone

Seven HMT inhibitors of the SET3 HMT G9a share a diaminoquinazoline backbone ^28-33^. Figure 5A shows pairs with >60% similarity and >50% delta activity at 1 µM in both species. For *Babesia divergens*, the side group on position 4 of the diaminoquinazoline scaffold seemed to have the most effect on compound activity (Figure 5B). A cyclohexylmethyl-4-piperidylamine side group (UNC0631) showed the highest activity, while the EC50 value increased 5-6 times when changing to a cyclohexyl-4-piperidylamine (UNC0646) or 1-benzyl-4-piperidylamine (BIX01294) side group. Substituting the ring structure with an isopropyl group (UNC0638 and UNC0642) further decreased the remaining activity by half. Activity is completely lost when the side group consists of a lone 4-piperidylamine (UNC0224).

Previous SAR studies of diaminoquinazolines methyltransferase inhibitors have been performed for *P. falciparum*, ^34,35^. We confirmed that substituting the 1-benzyl-4-piperidylamine of BIX01294 on position 4 of the diaminoquinazoline scaffold with a cyclohexylmethyl-4-piperidylamine (UNC0631) or a cyclohexyl-4-piperidylamine (UNC0646) reduced the activity against *P. falciparum* (**Supplemental** Figure 5). However, in disagreement with previous findings, the substitution with a 1-isopropyl-4-piperidylamine (UNC0642 and UNC0638) did not impact the activity against *P. falciparum*.

It was previously reported that a lysine mimetic side group on position 7 of the diaminoquinazoline scaffold lacks activity against *P. falciparum*, despite exhibiting potent activity against G9a due to interactions in the lysine binding channel of this enzyme. We confirmed that a dimethylpropylamine side group (UNC0224) loses activity as previously reported, but interestingly, we found the lysine mimetic side groups 1-propylpyrrolidine (UNC0642 and UNC0638) and 1-propylpiperidine (UNC0631 and UNC0646) retained most of their activity. This suggests that compounds with a lysine mimetic side group on position 7 of the diaminoquinazoline scaffold can inhibit *P. falciparum* only if combined with certain side groups on position 2 and 4 of the scaffold.

Three of the diaminoquinazoline compounds have a high differential activity against both species, showing 17-24 times more activity against *P. falciparum* (Figure 5C). As two of these compounds (UNC0642 and UNC0638) share a 1-isopropyl-4-piperidylamine group on position 4 and a 1-propylpyrrolidine group on position 7, these might contribute to this difference in activity. As the closest related *P. falciparum* HMT to HsG9a is PfSET3, it is possible that the binding site on the HMT enzyme for these three compounds is more divergent from HsG9a in BdSET3 than in PfSET3.

Overall, we show that epigenetic enzymes may be a promising novel target in *Babesia divergens*. Our library of 324 epigenetic inhibitors includes 19 pairs of compounds with high delta activity despite high structural similarity, which provide insight into the structural features that confer activity against *P. falciparum* and *B. divergens*. Multiple diaminoquinazoline backbone HMT inhibitors show highly active against both species tested, with UNC0631 displaying a low nanomolar range EC50 value against both *P. falciparum* and *B. divergens*, while UNC0224 is inactive against both species despite minor structural differences with the active diaminoquinazoline compounds.

## Supporting information

Supplementary Data Set 1

Supplementary Data Set 2

## ACKNOWLEDGEMENTS

We thank the High Throughput and Spectroscopy Resource Center at Rockefeller University for technical assistance, and Elisabeth Martinez (UT Southwestern) for additional JIB-04 inhibitor. We also thank Laura Kirkman for providing the *B. divergens* strain and valuable feedback on the manuscript. This work was supported by a Bohmfalk Charitable Trust Research Grant and NIH 1R01AI141965 and 1R01AI138499 to BK, and a Belgian American Educational Foundation post-doctoral fellowship to LV.

## SUPPLEMENTARY FIGURE LEGENDS

**Figure S1.**
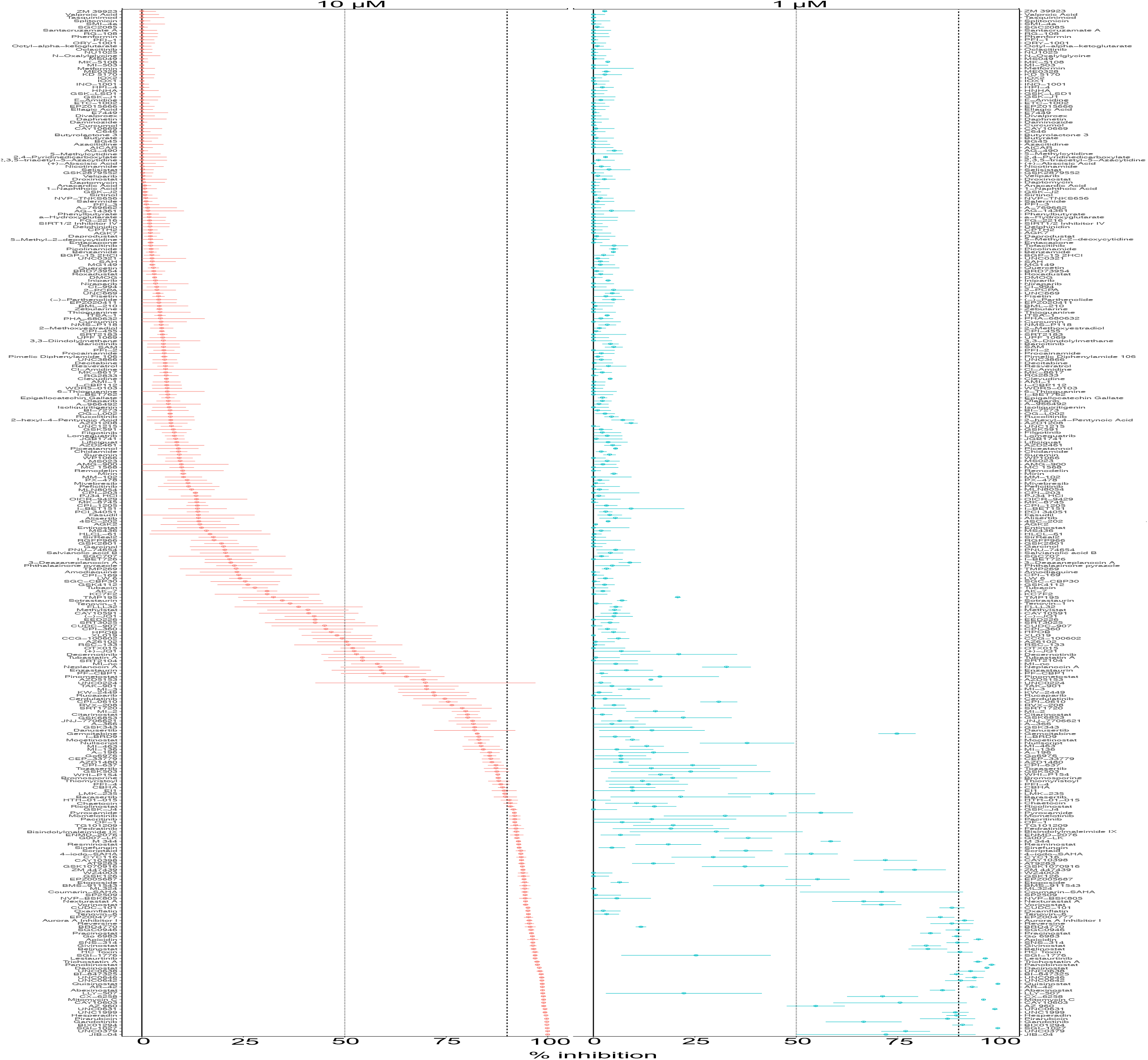
Mean Percent Inhibition of all 324 compounds against *B. divergens* at 10 μM and 1 μM. The dotted and dashed lines indicate 50% and 90% inhibition, respectively. Compounds are ordered by increasing activity at 10 μM. Error bars are standard error of n=3.

**Figure S2.**
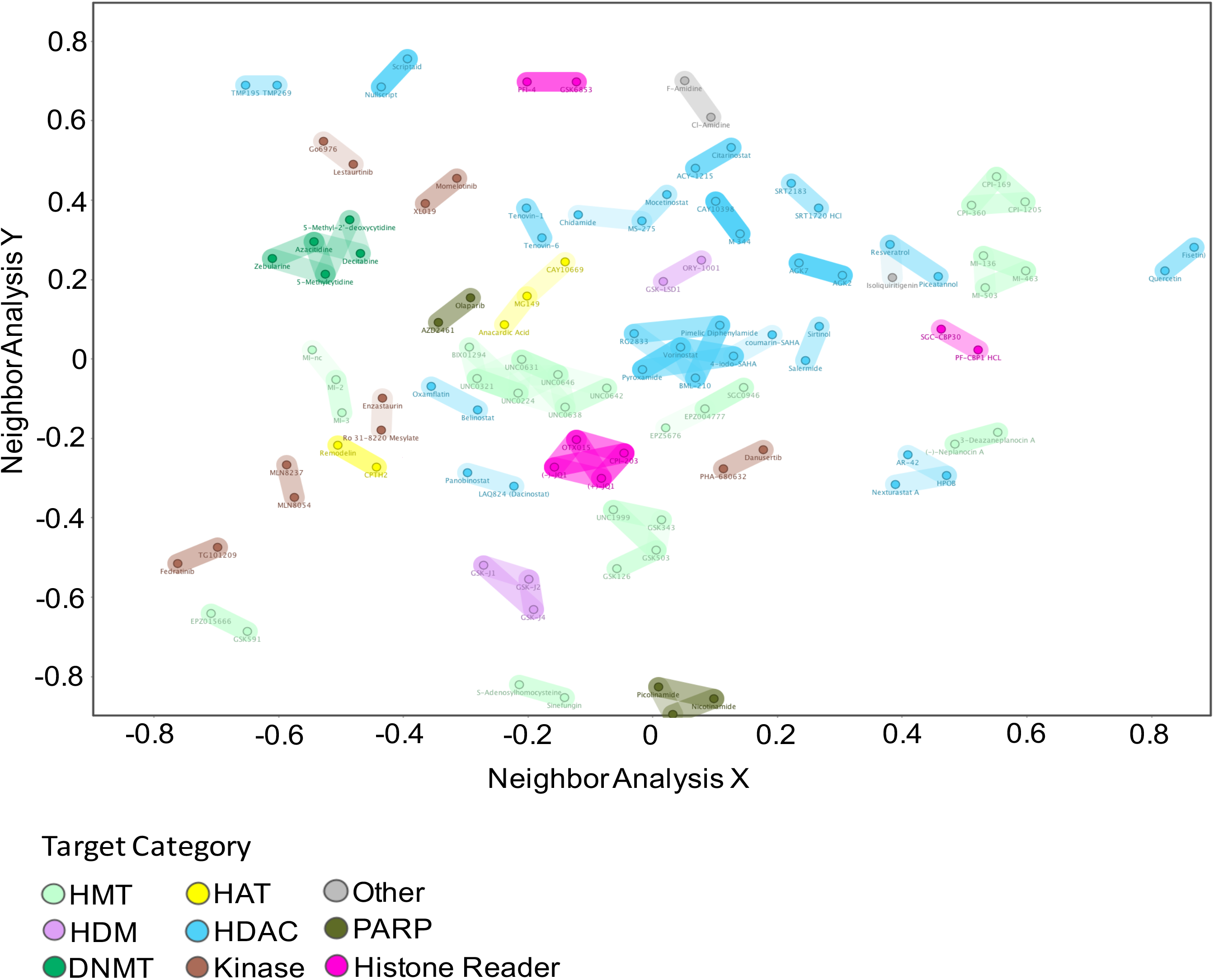
Structural feature similarity landscape. Compounds with >80% structural similarity were grouped in Datawarrior (SkelSphere). Color indicates reported epigenetic process targeted in higher eukaryotes.

**Figure S3.**
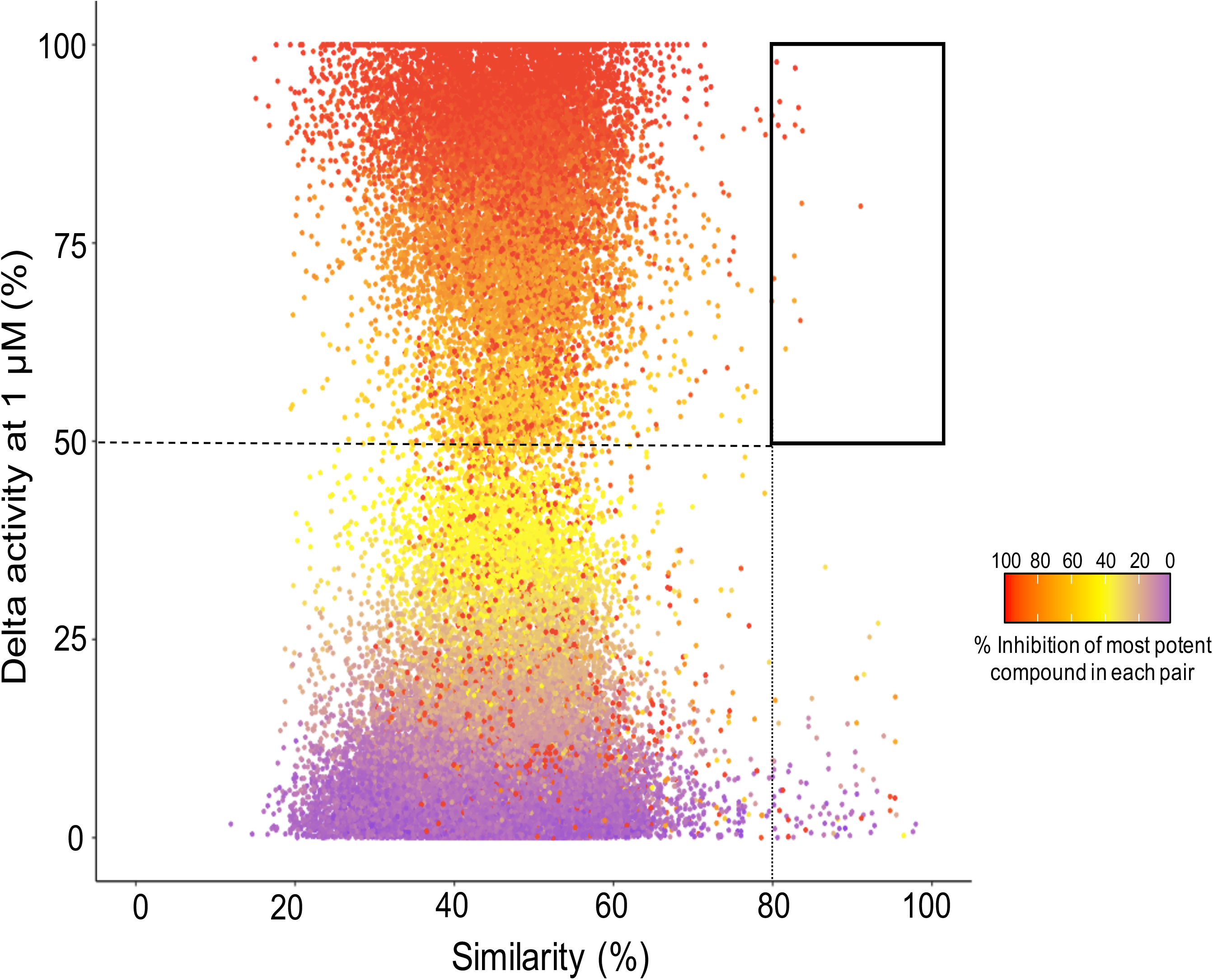
Activity cliff analysis for *Babesia divergens* at 1 μM. Scatterplot with each dot representing a pair of compounds in the library. Compound pairs of interest have ≥50% delta activity (dashed line) and ≥80% structural similarity (dotted line).

**Figure S4.**
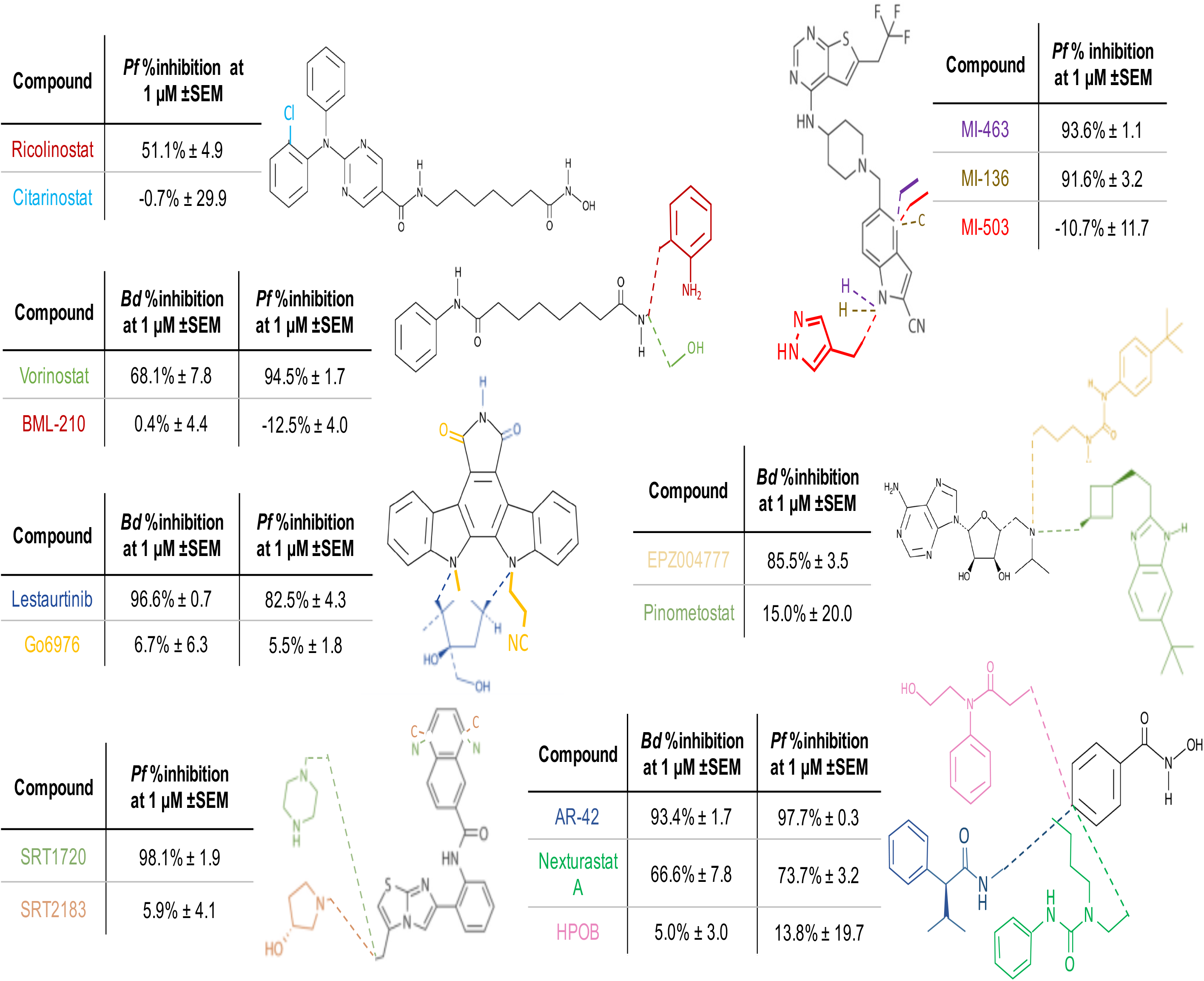
Structural representation of the remaining activity cliff pairs that are displayed in figure 4A.

**Figure S5.**
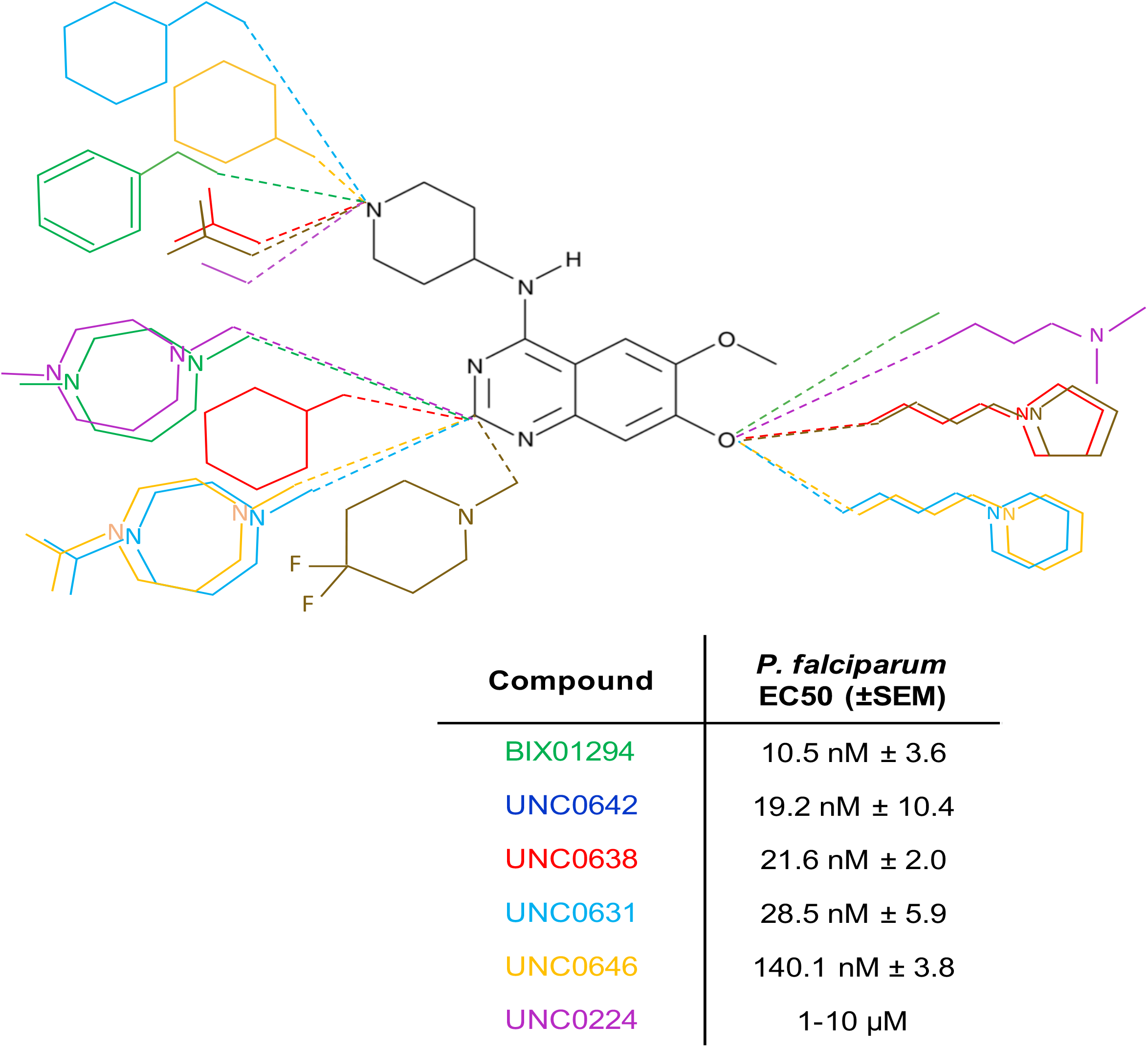
Activity of HMT inhibitors with a diaminoquinazoline backbone against *P. falciparum*.

**Table ST1.** EC50 Activity of epigenetic inhibitors tested grouped by target category. Percentage of active compounds is indicated in brackets (n=2-3).

| Target Class | Compounds | >50% inhibition at 10 μM | | >50% inhibition at 1 μM | |
| --- | --- | --- | --- | --- | --- |
|  |  | *B. divergens* | *P. falciparum* | *B. divergens* | *P. falciparum* |
| Histone Acetylation | 10 | 0 (0%) | 1 (10%) | 0 (0%) | 0 (0%) |
| Histone Deacetylation | 85 | 34 (40%) | 43 (51%) | 20 (24%) | 25 (29%) |
| Histone Methylation | 51 | 28 (55%) | 32 (63%) | 10 (20%) | 11 (22%) |
| Histone Demethylation | 18 | 4 (22%) | 9 (50%) | 1 (6%) | 1 (6%) |
| Histone Phosphorylation | 66 | 38 (58%) | 41 (62%) | 11 (17%) | 13 (20%) |
| Histone PARPylation | 22 | 3 (14%) | 5 (23%) | 0 (0%) | 0 (0%) |
| Histone Reader Domains | 28 | 12 (43%) | 6 (21%) | 0 (0%) | 0 (0%) |
| DNA Methylation | 14 | 2 (14%) | 3 (21%) | 2 (14%) | 1 (7%) |
| Other | 30 | 4 (13%) | 6 (20%) | 2 (7%) | 3 (10%) |
| **Total** | **324** | **125 (39%)** | **146 (45%)** | **46 (14%)** | **54 (17%)** |

**Table ST2.** EC90 Activity of epigenetic inhibitors tested grouped by target category. Percentage of active compounds is indicated in brackets (n=2-3).

| Target Class | Compounds | >90% inhibition at 10 μM | | >90% inhibition at 1 μM | |
| --- | --- | --- | --- | --- | --- |
|  |  | *B. divergens* | *P. falciparum* | *B. divergens* | *P. falciparum* |
| Histone Acetylation | 10 | 0 (0%) | 0 (0%) | 0 (0%) | 0 (0%) |
| Histone Deacetylation | 85 | 25 (29%) | 34 (40%) | 7 (8%) | 15 (18%) |
| Histone Methylation | 51 | 15 (29%) | 19 (37%) | 6 (12%) | 9 (18%) |
| Histone Demethylation | 18 | 4 (22%) | 4 (22%) | 0 (0%) | 1 (6%) |
| Histone Phosphorylation | 66 | 25 (38%) | 25 (38%) | 2 (3%) | 5 (8%) |
| Histone PARPylation | 22 | 1 (5%) | 4 (18%) | 0 (0%) | 0 (0%) |
| Histone Reader Domains | 28 | 1 (4%) | 1 (4%) | 0 (0%) | 0 (0%) |
| DNA Methylation | 14 | 1 (7%) | 1 (7%) | 1 (7%) | 1 (7%) |
| Other | 30 | 3 (10%) | 4 (13%) | 1 (3%) | 1 (3%) |
| **Total** | **324** | **75 (23%)** | **92 (28%)** | **17 (5%)** | **32 (10%)** |

**Supplemental Dataset 1**: Gene identifiers of epigenetic writer and reader enzyme orthologs in Figure 1.

**Supplemental Dataset 2**: Mean percent inhibition of all compounds against B. divergens at 10 μM and 1 μM.

